# SDF-1 Bound Heparin Nanoparticles Recruit Progenitor Cells for Their Differentiation and Promotion of Angiogenesis After Stroke

**DOI:** 10.1101/2023.07.05.547800

**Authors:** Katrina L. Wilson, Lauren A. Onweller, Neica I. Joseph, Jennifer David-Bercholz, Nicole J. Darling, Tatiana Segura

**Affiliations:** Department of Biomedical Engineering, Duke University, Durham NC 27708-0281, USA; Department of Anesthesiology, Duke University, Durham NC, 27708-0281, USA; Department of Chemical and Biomolecular Engineering, University of California Los Angeles, Los Angeles, CA 90095, USA; Department of Neurology, Duke University, Durham, NC, 27708-0281 USA; Department of Dermatology, Duke University, Durham, NC, 27708-0281 USA

**Keywords:** MAP, hydrogel, microgel/microparticle, neural stem cells, cryogels, stroke, SDF-1

## Abstract

Angiogenesis after stroke is correlated with enhanced tissue repair and functional outcomes. The existing body of research in biomaterials for stroke focuses on hydrogels for the delivery of stem cells, growth factors, or small molecules or drugs. Despite the ability of hydrogels to enhance all these delivery methods, no material has significantly regrown vasculature within the translatable timeline of days to weeks after stroke. Here we developed 2 novel biomaterials for tissue regeneration after stroke, a highly porous granular hydrogel termed Cryo microgels, and heparin-norbornene nanoparticles with covalently bound SDF-1α. The combination of these materials resulted in fully revascularized vessels throughout the stroke core in only 10 days, as well as increased neural progenitor cell migration and maintenance and increased neurons.

## 1. Introduction

Angiogenesis plays a major role in regeneration after ischemic stroke in the brain, as healthy vasculature is essential for repairing and maintaining cells after thus type of injury. This complex physiological event relies on the balance of cellular responses with growth and transcription factors.^[1]^ Previous work in the biomaterials field has attempted to mimic or enhance natural tissue repair through the addition of growth factors, incorporation of stem cells, or changing chemical and physical properties of the biomaterials. Work in our lab has leveraged changing the structural properties of hydrogels by designing a granular scaffold called microporous annealed particle (MAP) scaffolds to enhance the porosity.^[2]^ These MAP scaffolds caused an increase in tissue regeneration in the brain after stroke, compared to nonporous hydrogels.^[3, 4]^ It is possible the increased porosity of our material led to enhanced repair in the hydrogel and tissue surrounding the stroke, as it is well known that the microstructure of a hydrogel scaffold influences cell-material interactions, such as cell attachment and migration and protein adsorption or adsorption.^[5, 6]^

Alternative avenues for using biomaterial for tissue repair in the brain after stroke lie in the use of growth factors or small molecules and stem cell delivery. Both the delivery of neural progenitor cells (NPCs)^[7–11]^ and the use of growth factors^[12–16]^ have been implored for tissue repair in the brain after stroke. Growth factors and small molecules are important after injury because they are largely responsible for determining cell fate, recruiting cells, and directing mechanisms for repair.^[17–20]^ SDF-1α, for example, is a chemoattractant molecule secreted after stroke and known to recruit NPCs,^[21, 22]^ endothelial progenitor cells (EPCs),^[15, 23]^ and mesenchymal stem cells to the infarct region.^[24, 25]^ Interestingly, NPCs and angiogenesis have been linked through crosstalk signaling between the NPCs and EPCs.^[26–28]^ Both NPCs and EPCs have been observed to secrete growth factors and chemokines that aid one another in migration and proliferation, as well as protection against hypoxia after stroke.^[29]^ In a previous study, researchers found that while NPCs attempt to migrate towards the injury site after stroke, only about 20% of new neurons survive 6 weeks after ischemic injury.^[30]^ This has made targeting the recruitment and survival of NPCs within the stroke cavity difficult. A possible avenue is through the administration of SDF-1α. Treatment of stroke with SDF-1α upregulates the SDF-1/CXCR4 axis and enhances neurogenesis, angiogenesis, and neurological outcome.^[25]^ Undesirably, the most common form of administration is via direct injection, which severely limits its therapeutic use due to systemic adverse effects and poor transport of large molecule drugs into the brain.^[31]^ To circumvent these issues, biomaterials have been used to deliver growth factors and small molecules, mitigating otherwise adverse effects and promoting a longer shelf life and sustained release. ^[32–35]^ Notably, there is a substantial gap and need for developing a biomaterial platform for enhancing the recruitment of endogenous NPCs for angiogenesis purpose.

Herein we report 2 novel biomaterial components for tissue regeneration after stroke, a highly porous granular hydrogels termed Cryo microgels, and heparin-norbornene nanoparticles with covalently bound SDF-1α. Cryo scaffolds were first implored to enhance vascularization solely through the biomaterial’s physical properties. Initial results demonstrated an increase in vasculature with increased porosity. However, the vasculature did not penetrate the entire stroke core. The addition of SDF-1 was used to increase NPC migration and maintenance in hopes of promoting vessel formation through cell-cell mediated cross talk.^[36–38]^ When Cryo and SDF-1 were used together, we observe revascularization of the stroke infarct in only 10 days after injection, or 15 days after stroke, as well as NPC migration and neuronal marker labeling within our material. To our knowledge, this is the first demonstration of using cryo microgel scaffolds in the brain after stroke as well as covalently bound SDF-1α nanoparticles to completely revascularize the entire stroke core in only 10 days.

## 2. Results and Discussion

### 2.1 Creating Cryo Microgels Increases Porosity While Maintaining Storage Modulus

Previous work in our lab has demonstrated the injection of MAP scaffolds following stroke to attenuate astrocyte and microglia/macrophage reactivity.^[3, 4]^ These observations were first examined in comparison to nonporous hydrogels, which typically require cells to degrade the scaffold before laying down their own matrix. It was concluded that the enhanced porosity and structure of the MAP scaffold was responsible for the decrease in inflammatory response. As such, we believed that increasing the porosity further could aid in enhancing cellular infiltration, increasing diffusion of surrounding molecules that are naturally released after stroke, or act as a sponge for drug depot delivery. Therefore, we sought after possible mechanisms to increase MAP scaffold porosity. At first, the idea of changing microgel size to alter void space seemed natural. However, simulated data using LOVAMAP shows that changing microgel size does not alter void space.^[39]^ Instead, only by changing the packing of the microgels can we alter the percentage of void space within the scaffold. Additionally, recent publications from our lab using poly(ethyl glycol) microgels confirmed the LOVAMAP simulated data *in vitro*.^[40]^ Another recent publication in our lab also showed mechanisms to change the packing of our MAP scaffolds and demonstrated the enhanced effects this has on cell spreading *in vitro.*^[38]^ However, the controlled packing of the MAP scaffold in this mechanism only allowed for controlled particle fraction from 86%-98%, or void volume fraction of 2-14% in the MAP scaffold porosity. Therefore, if one wanted to study a MAP scaffold with greater than 24% void volume fraction, alternative methods would be required.

Historically increasing bulk hydrogel porosity could be achieved through electrospinning hydrogels, creating precast hydrogels, or by the formation of cryogels.^[41, 42]^ We adapted the process of making cryogels to our microgels in hopes to enhance porosity of the individual microgels and therefore overall MAP scaffolds. First HMPs were centrifuged at 4,500rpm for 5 min and excess buffer was removed with a kimwipe to “dry” the microgels. Then HMPs were resuspended in 200 mM TrisHCl, 150 mM NaCl, 20 mM CaCl2 buffer overnight. The microgels were then dried using the same protocol and flash frozen in liquid nitrogen, resulting in ice crystal formation, and lyophilized to lock in the void space formed through flash freezing, resulting in the formation of cryogels (**Figure 1a**). The lyophilization allowed sublimation to occur, leaving the microgels, now Cryo microgels, with an interconnected microporous structure as compared to the nanoporous structure of the original microgels. To assess porosity, microgels and cryogels were resuspended in 7,000 kDa fluorescein-labeled dextran solutions and imaged using confocal microscopy, and then visualized using Nikon rendering to compare pores between and inside of gels, labeled with the dextran (**Figure 1b**). The dextran size was chosen, as it readily diffused throughout open pores but did not penetrate nanoporous microgels or nanoporous structure of the cryogels. Microparticles from microgels and cryogels were visualized after confocal imaging using IMARIS software to better understand the morphology differences between pores in each condition The void fractions of the MAP scaffolds and Cryo scaffolds (**Figure 1c**) were shown to be 0.287 ± 0.02 and 0.4407 ± 0.015, respectively (**Figure 1d**). Further material characterization was performed to measure if increased porosity within the cryogel scaffolds would change the moduli. Both Young’s and Storage moduli were tested (**Figure 1d-g**). No statistical difference was observed between scaffolds formed at an R7, with MAP scaffolds having a storage modulus of 373.64 Pa and Cryo scaffolds having a storage modulus of 393.93 Pa.

**Figure 1.**
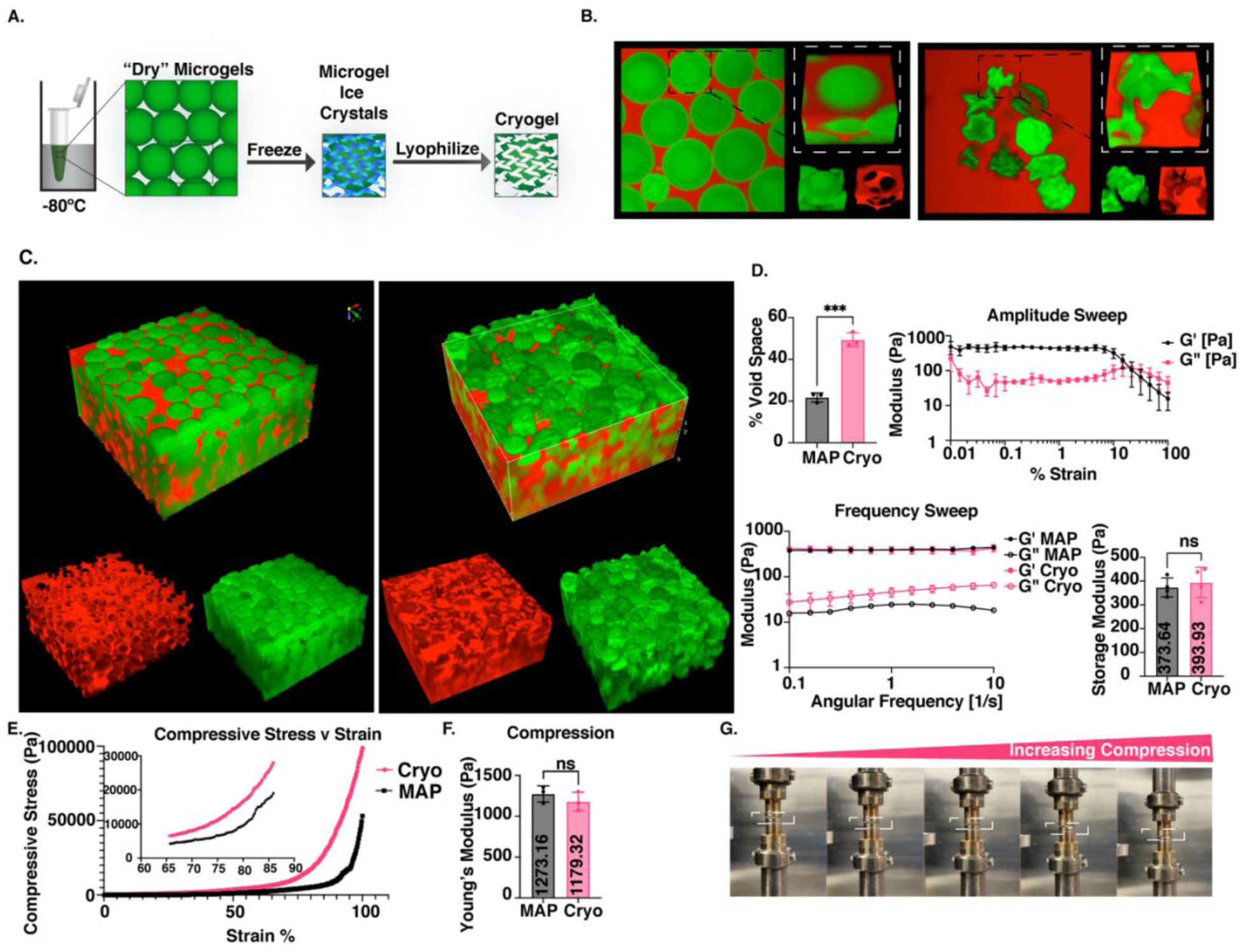
Formation of cryo microgels and material characterization. A. Schematic showing how cryogels are made. B. Confocal images and Nikon volume renderings of microgels compared to cryogels. C. IMARIS surface renderings of MAP scaffolds compared to Cryo scaffolds. D. Quantification of percent void space from IMARIS renderings, Amplitude sweep of Cryo Scaffold, Frequency sweep of MAP and Cryo Scaffolds, Storage modulus of MAP scaffold verses Cryo scaffold. E. Compressive stress over strain of Cryo and MAP scaffolds. F. Calculated Young’s Modulus from compressing tests. G. Compressive test images over increasing strain.

We were able to confirm that MAP microgels and cryogels had visually and quantitatively different microarchitectures, with significantly more void space in cryogels. At the same time, the material properties on the scaffold level scale were consistent with each other and had no statistically significant differences. These results were consistent with the idea that the composition of the scaffolds are similar on the scaffold scale, but on the microparticle scale the gels differ. Previous studies in the field have shown that cells sense forces and signals on the micro scale, giving further motivation to see how cells respond to microarchitecture changes between MAP microgel and cryogel conditions in later experiments.^[43]^ For this reason, we decided to move forward with making and testing cryogels to study how cells would respond to Cryo scaffolds composed of cryogel microparticles compared to MAP, and investigate if porosity could enhance angiogenesis in the brain after stroke.

### 2.2 Enhanced Porosity Increases Vessel Infiltration but Has No Change in GFAP Scar Thickness or Infiltration compared to MAP Scaffolds

Initial animal testing investigated the use of MAP versus Cryo gel injected into mice 5 days after a photothrombotic stroke (PT) stroke and sacrificed on day 15 (**Figure 2a**). Both hydrogels were modified to contain RGD peptides only. Tissue was collected 10 days after injection, cryosection, IHC stained, and imaged for analysis with a 5×5 tile grid at 20x magnification and oriented with the stroke in the top of the image and peri infarct towards the sides and bottom of the image (**Figure 2b**). To compare initial immune responses, glial fibrillary acidic protein (GFAP, astrocyte marker) was stained for, as well as ionized calcium binding adaptor molecule 1 (Iba1, microglia/macrophage marker) (**Figure 2c**). Further stains with neurofilament 200 (NF200, neuron marker) and glucose transporter 1 (Glut1, vasculature marker) were also performed to measure tissue regeneration (**Figure 2c**). Analysis revealed no significant difference between MAP scaffolds and Cryo scaffold GFAP+ and Iba1+ groups (**Figure 2d**). Scar thickness was quantified through measuring the GFAP+ distance in the border of the stroke area, making sure not to quantify the corpus callosum (**Figure 2d**). Glut1+ cells revealed a significant increase in vessel infiltration in the infarct of Cryo treated mice 10 days after gel injection compared to MAP (**Figure 2d**). Interestingly, there was also a significant difference in neurons between MAP and Cryo scaffolds (**Figure 2d**), where Cryo had a significant decrease in positive area of NF200 compared to MAP. While vessel increase was significant, the vessels did not penetrate past 200 µm into the infarct (**Figure 2d**), therefore we sought to incorporate the delivery of SDF-1 nH in hopes to enhance the recruitment of NPCs and vessel formation via cell-cell mediate crosstalk.

**Figure 2:**
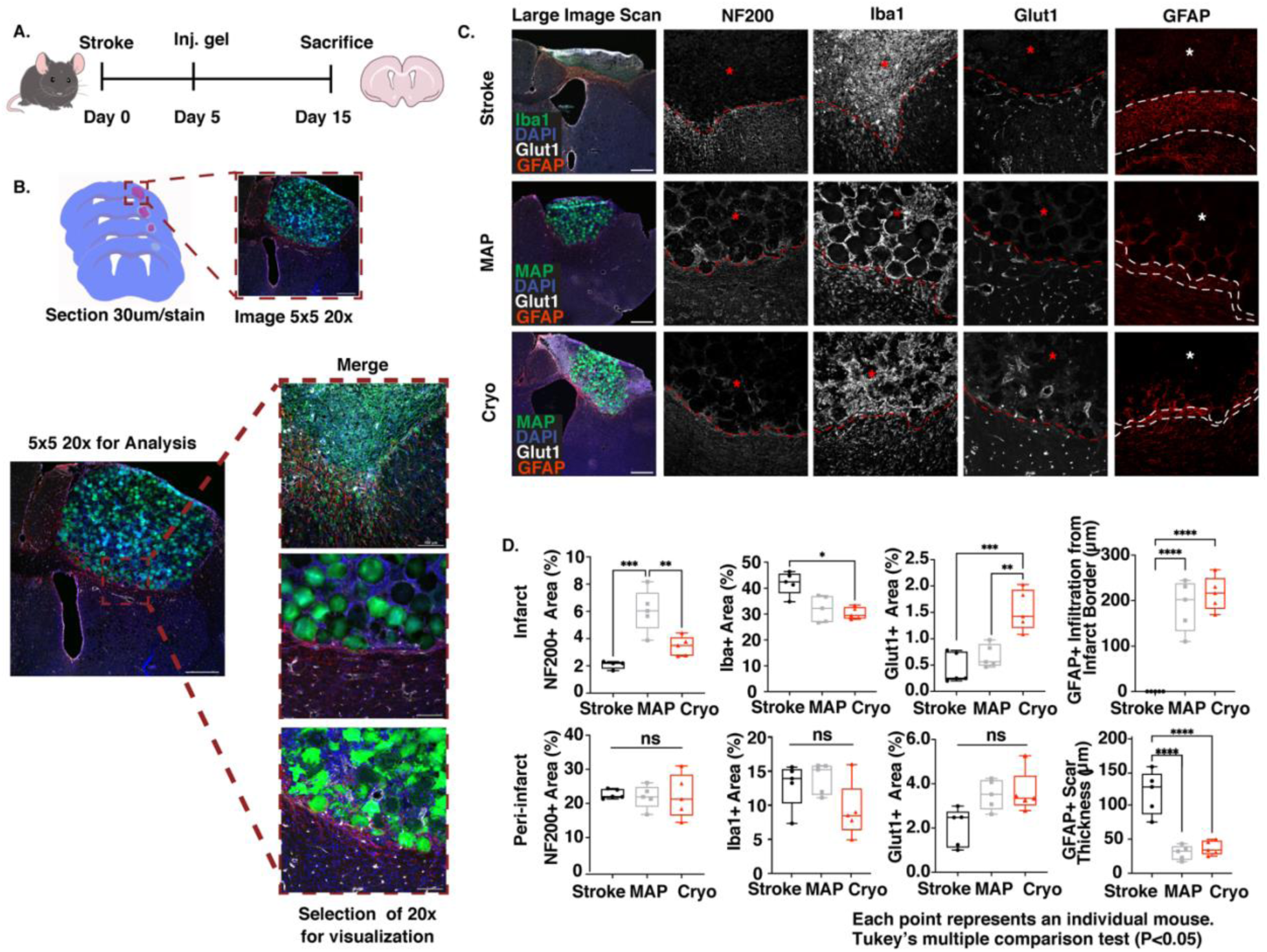
Effects of porosity on Glut1+ vessel infiltration 10 days after stroke. A) Depicts *in vivo* experimental overview of photothrombotic stroke on Day 0, gel injections on Day 5, sacrificing on Day 15. B) Depicts imaging of ROI brain sections and orientation of 20x representative images in C. For each image, the fourth section of the slide was imaged with a 5×5 tile grid at 20x magnification and oriented with the stroke in the top of the image and peri infarct towards the bottom of the image. Representative stained for Vessels (Glut1+) in white, astrocytes (GFAP+) in red, MAP in green, and nuclei (DAPI+) in blue. C) Representative images from stroke only, MAP, and Cryo scaffold conditions are shown stained for neurons (NF200+), microglia/macrophages (Iba1+), Vessels (Glut1+), and astrocytes (GFAP+). Dotted line indicates the region between peri-infarct and infarct (with an asterisk). D) Analysis of infarct and peri-infarct areas. (D) The percent positive area of neurons (NF200), microglia/macrophages (Iba1), vessels (Glut1), and astrocytes (GFAP) was measured across conditions in the stroke infarct area as well as the peri infarct area next to the stroke. Scale bars represent 500 microns (N=5).

### 2.3 Heparin-Norbornene Nanoparticle Synthesis Enables for Efficient and Permeant Tethering of SDF-1 in Cryo MAP Scaffolds

Previous work in our lab created heparin nanoparticles (nH) that would covalently bind growth factors through light sensitive free-radical reactions.^[13]^ However, we sought to modify this method to enable tethering of the nanoparticles to the surface of microgels in a robust, non-light sensitive method. Like previous work,^[44]^ this was achieved through modifying the heparin sulfate to have norbornene functional handles for both creating the nH and future tethering of SDF-1 or other growth factors and binding to MAP scaffolds via NB-Tet click chemistry (**Figure 3a**). Modification was determined via H’NMR (**Figure S1**), where 62% modification was sufficient for both creating nH and post modification. The heparin-norbornene polymer was stored at −20°C until its use for nH production. Nanoparticles were created via sonication of the heparin-norbornene polymer in an inverse emulsion. Once purified, scanning electron microscopy (SEM) (**Figure 3b**) and dynamic light scattering (DLS) (**Figure 3c**) was performed to determine nanoparticle size distribution. Some bimodal or spreading was initially observed (data not shown) but with the use of a 0.2 μm filter we could obtain relatively monodispersed (PDI of 0.21 or less) nH, averaging 100 nm. These nH were used to covalently bind SDF-1 growth factor to the norbornene groups with LAP and UV light on ice. After 2 min of 20mW UV light in 10 mM LAP 1x PBS, the SDF-1 nH were purified through centrifugation and the flowthrough was measured for any unbound SDF-1 to determine the binding efficiency (**Figure 3d**). Minimal SDF-1 growth factor, about 15%, would not bind to the nH. This concentration was consistent each time and resulted in a ratio of about 9.90 ng SDF-1:nH. To test the validity of the of the SDF-1 nH binding to our scaffold, a release assay was performed in which 200 ng of SDF-1 was mixed with HA-NB microgels and crosslinked with 4-arm PEG-Tet to create a MAP scaffold. The scaffold was left in 1x PBS at 37°C for 11 days. PBS was removed and replaced at each time point indicated and then an ELISA was performed to test for SDF-1 release. No release was observed (**Figure 3e**) indicating that the growth factor remained bound to the nH and the nH remained bound to our scaffold.

**Figure 3:**
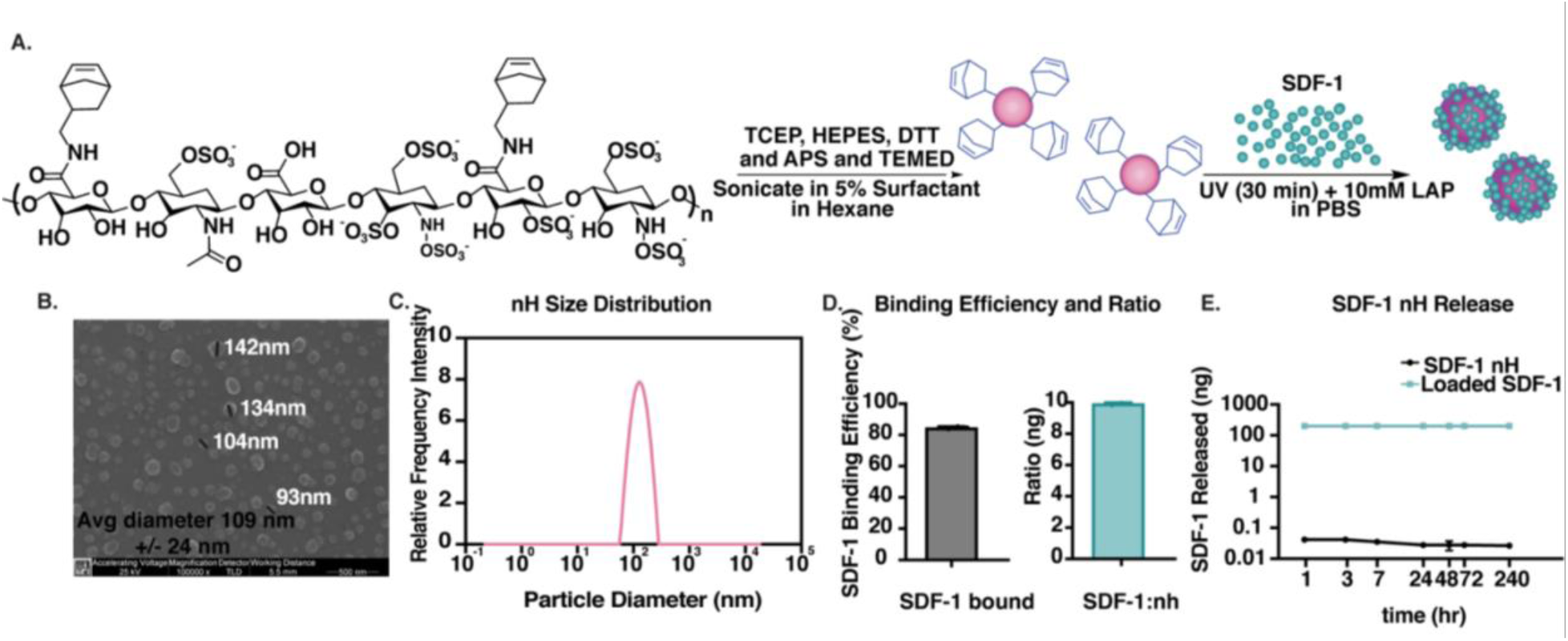
Synthesis of Heparin Nanoparticles for Delivery of SDF-1. A) Chemical schematic of synthesizing heparin-norbornene nanoparticles (nH) from heparin modified with norbornene and covalent binding of SDF-1 soluble growth factor to nH. B) SEM image of gold coated nH. C) DLS data of nH, PDI = 0.23. D) ELISA data of measured SDF-1 bound to nH and ratio of SDF-1 to nHs. E) Release assay of SDF-1 nH in MAP scaffolds. Blue line indicates the amount of SDF-1 loaded in the scaffold.

### 2.4 SDF-1 Nanoparticles Increase CXCR4 Expression 6 Hours After Exposure and Increases Cell Spreading *In Vitro*

SDF-1 growth factor is known to mediate NPC migration and interaction via the CXCR4 pathway.^[25]^ We hoped to use this mechanism to recruit NPCs after stroke. Previous methods in the field look to add SDF-1 as an unbound molecule to allow for gradients emitting from the site of delivery.^[45–47]^ Alternatively, we hoped that by binding the SDF-1 covalently to the nH we would promote endogenous migrating NPCs to the stroke region to enter and be sustained within our material. We theorized that NPC maintenance would allow for cell-mediated crosstalk with ECs to enhance vessel formation or NPC differentiation into neurons. We first went on to determine the concertation of SDF-1 that would activate NPCs via CXCR4 signaling. This was done by indirectly measuring the CXCR4 activation via RT-PCR of 2D cultured NPCs after their exposure to various soluble concentrations of SDF-1 (**Figure 4a**). Interestingly, we found there to be an optimal concentration of SDF-1. Exposure from 100 to 200 ng of SDF-1 led to an increase in CXCR4 gene expression at 6 hours but increasing the amount of SDF-1 past 200 ng did not result in neither higher gene expression nor longer expression, but instead had no change in CXCR4 gene expression (**Figure 4a, Sup fig X**). There was also an observed bimodal response to the 200 ng SDF-1 exposure, where at 6 hours we saw the highest increase, a decrease at 24 hours and then another increase at 48 hr (**Sup Fig S2**). We concluded that 200 ng of SDF-1 was sufficient to increase CXCR4 expression over 2-fold compared to housekeeping genes. However, as expected, this activation was not sustained past 6 hours, possibly due to SDF-1 cellular uptake or degradation over time.

**Figure 4:**
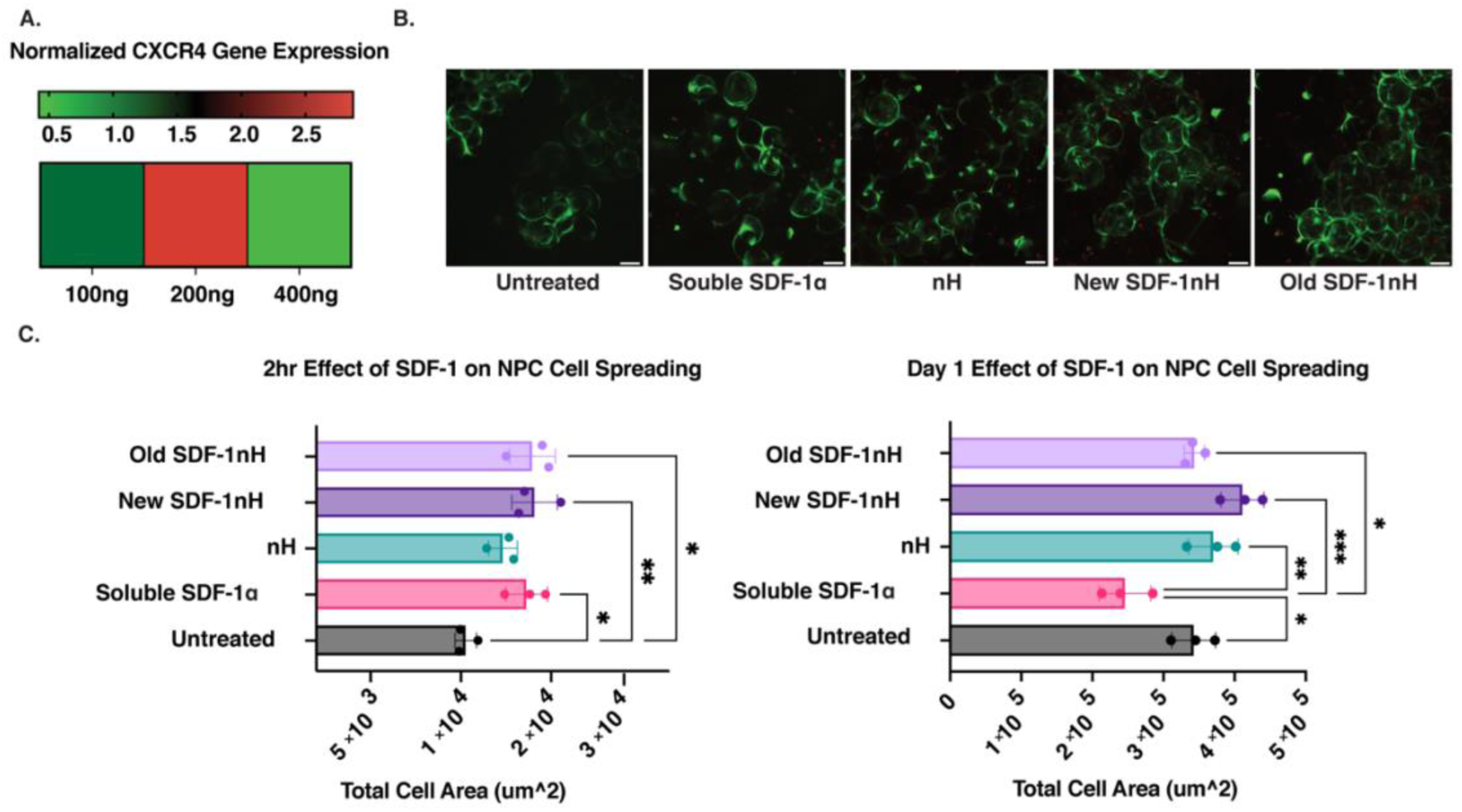
SDF-1 Effects on NPCs *in vitro.* A) RT-PCR measuring CXCR4 expression after NPC exposure to concentrations of 100 ng, 200 ng, and 400 ng of soluble SDF-1. B) Representative live/dead immunofluorescent images of NPCs spreading in MAP Scaffolds at D4 stained with CalceinAM. Scale bars represent 100 microns (N=3). C) Quantification of NPC spreading after 2 hours and 1 day of exposure to nH soluble SDF-1, bound and unbound SDF-1 nH at 200ng SDF-1.

Since covalent binding of growth factors can have detrimental effects on their function, we wanted to test if the bound SDF-1 was still active. We did this by exposing NPCs within our MAP scaffold to 200 ng of bound SDF-1 nH, 200 ng soluble SDF-1, or nH alone, as well as an untreated control. Due to difficulties in purifying NPCs from the hydrogel for RT-PCR, we turned to measuring cell spreading to determine if the SDF-1 bound to the nH was still active. Measurements and/or images of cell spreading over the course of 4 days after exposure to treatment showed significant increases in cell spreading in the groups treated with SDF-1 nH within the first day compared to soluble SDF-1 and CalceinAM live/dead staining on day 4 showed high viability and spreading in SDF-1 nH conditions (**Figure 4b & c**). Since we wanted to validate the shelf life of the SDF-1 once bound to the NPs, we kept SDF-1 nH for 2 weeks at 4°C. This “old” SDF-1 nH still showed a significant increase in cell spreading compared to soluble SDF-1, indicating that binding the SDF-1 to our nH retained the SDF-1 bioactivity. Interestingly, on Day 1, nH only also showed an increase in cell spreading, though not as significant as the new SDF-1 nH. We believe that the presence of the naked particles could sequester growth factors naturally secreted by NPCs, allowing them to have sustained effect that may slightly enhance their cell growth.

### 2.5 SDF-1 Incorporation recruits SOX2+ NPCs and results in Tuj1+ Neuronal Differentiation

Before injection, Cryo microgels were post-modified with laminin derived IKVAV, YIGSR, and RGD peptides at concentrations that dictated NPC differentiation into Tuj1+ cells in previous work *in vitro.*^[44]^ Therefore, we hypothesized that NPCs found entering our scaffolds would differentiate into Tuj1+ neuronal cells. Following successful *in vitro* experimentation, we proceeded to incorporate the SDF-1 nH into our Cryo scaffolds prior to injection, at a 200 ng/6 μl concertation. Nestin-Cre/Ai9 mice were bred in attempts to lineage trace NPCs through the labelling of nestin positive cells and see if they differentiated into GFAP+, Tuj1+, or other neuronal cell types within our material. Unfortunately, in the system we used, Cre-mediated recombination was induced at birth, making it difficult to trace new progenitors formed close in time to material injection and after the neonatal state. The system resulted in bright mCherry signal in cells that were Nestin+ at some point in their development such as neural stem and progenitor cells, glia precursor cells, mesenchymal stromal cells, and endothelial progenitor cells.^[48]^ Interestingly, Nestin+ cells in the stroke core die, leaving the infarct mCherry negative, which allowed us to track Nestin+ cells migrating to the stroke core, which we believe to be progenitor and stem-like cells. For future studies, a CreER system could be used to induce Cre-mediated recombination with tamoxifen injections to ensure that only nestin positive cells from the time of interest would be labeled. Brain samples were collected 10 days after injection, or 15 days post stroke, as performed earlier (**Figure 5a**). Samples section were 30 μm and stained with SOX2, a known NPC marker, which turns off after differentiation, to investigate NPC cell migration into our scaffolds (**Figure 5b&c**).^[49]^ We observed SOX2+ cells in both Cryo scaffold and Cryo scaffold SDF-1nH groups, with a significant increase in the percent of SOX2+ cells in the Cryo scaffold SDF-1nH treated group (**Figure 5b&c**). While NPCs are known to migrate to the stroke core after injury, few survive the migration or are maintained at the stroke core.^[30]^ Here we see both the appearance of NPC migration from the SVZ and maintained NPC survival within our material 10 days post injection at a significantly higher percentage than Cryo scaffold only (**Figure 5b&c**).

**Figure 5:**
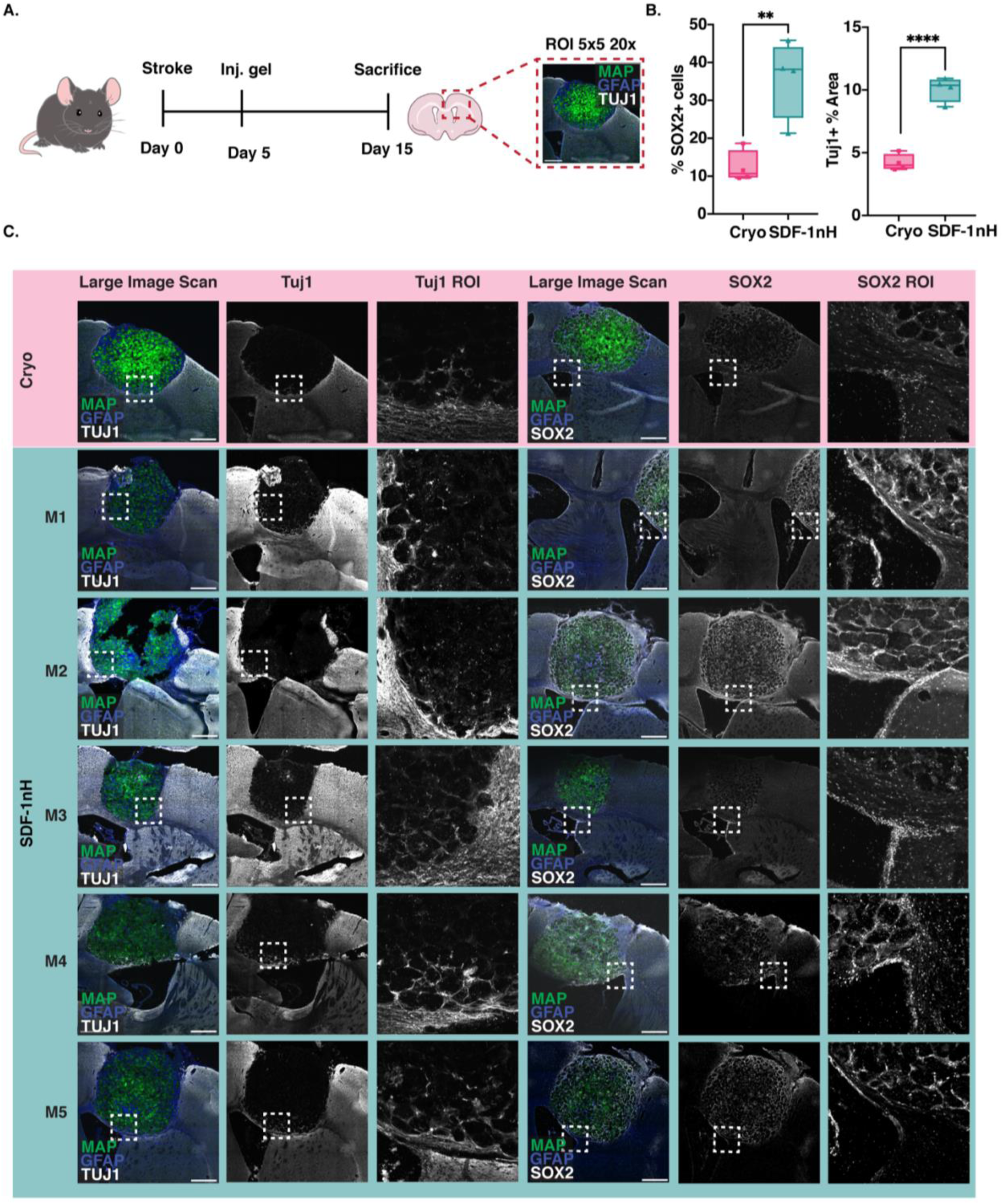
SDF-1 nanoparticles recruit SOX2+ and Tuj1+ cells inside SDF-1 Cryo Scaffolds. A) Experimental timeline in vivo for stroked mice injected with MAP and schematic explaining imaging of fixed and sectioned brains. B) Quantification of colocalization of SOX2+ cells and DAPI, and Tuj1+ percent area. Quantification was performed on large image scans (5×5, 20x). Cryo scaffolds (green), Tuj1+ cells (white), SOX2+ cells (White), Nestin+ cells (red), DAPI (blue). C) Large image scans and 20x magnified images for mice treated with Cryo scaffolds and Cryo scaffold with SDF-1 nH, stained for embryonic stem cell marker SOX2 (white), neuron marker Tuj1 (white), nestin (red), nuclei marker DAPI (blue), and labeled Cryo scaffolds (green). Scale bars represent 500 microns (N=5).

To determine if the enhanced NPC population we observed were differentiating into neuronal cells, we stained for neuron specific class II beta tubulin, or Tuj1, a marker known to be present starting in less mature neurons compared to other markers such as MAP2 (**Figure 5c**).^[50]^ Both Cryo scaffold and Cryo scaffold SDF-1nH showed the presence of Tuj1+ cells within the scaffold after 10 days, however, there was a significant increase in percent area of Tuj1+ cells in the Cryo scaffold SDF-1 nH condition compared to Cryo scaffolds only (**Figure 5b&c**). This is the first time we have reported observing Tuj1+ cells in our scaffolds 10 days post injection.

### 2.6 Delivery of SDF-1 in Cryo MAP Scaffolds Results in Significantly Increased Perfused Vasculature

Our main goal was to observe NPC-EC interaction and see if increasing the recruitment of NPCs would result in an increase of vessel formation. Previous Glut1+ stains showed vasculature in the Cryo scaffold model (**Figure 2c**) but did not indicate if the vasculature was connected to current blood vessels. To determine if this was the case, mice were perfused with tomato-lectin (TL) before sample collection. TL circulated in an awake mouse for 10 min prior to perfusions. Large image scans of 5×5, 20x images allowed us to capture the entire infarct for analysis and vasculature was traced to measure co-localization of Nestin+ cells that we hypothesized to be NPCs or ECs (**Figure 6a**). As hoped, there was a significantly increased percent positive area of vasculature in the SDF-1 nH Cryo scaffold group compared to Cryo scaffold alone group (**Figure 6b & c**). To our knowledge, this is the fastest revascularization of stroke.

**Figure 6:**
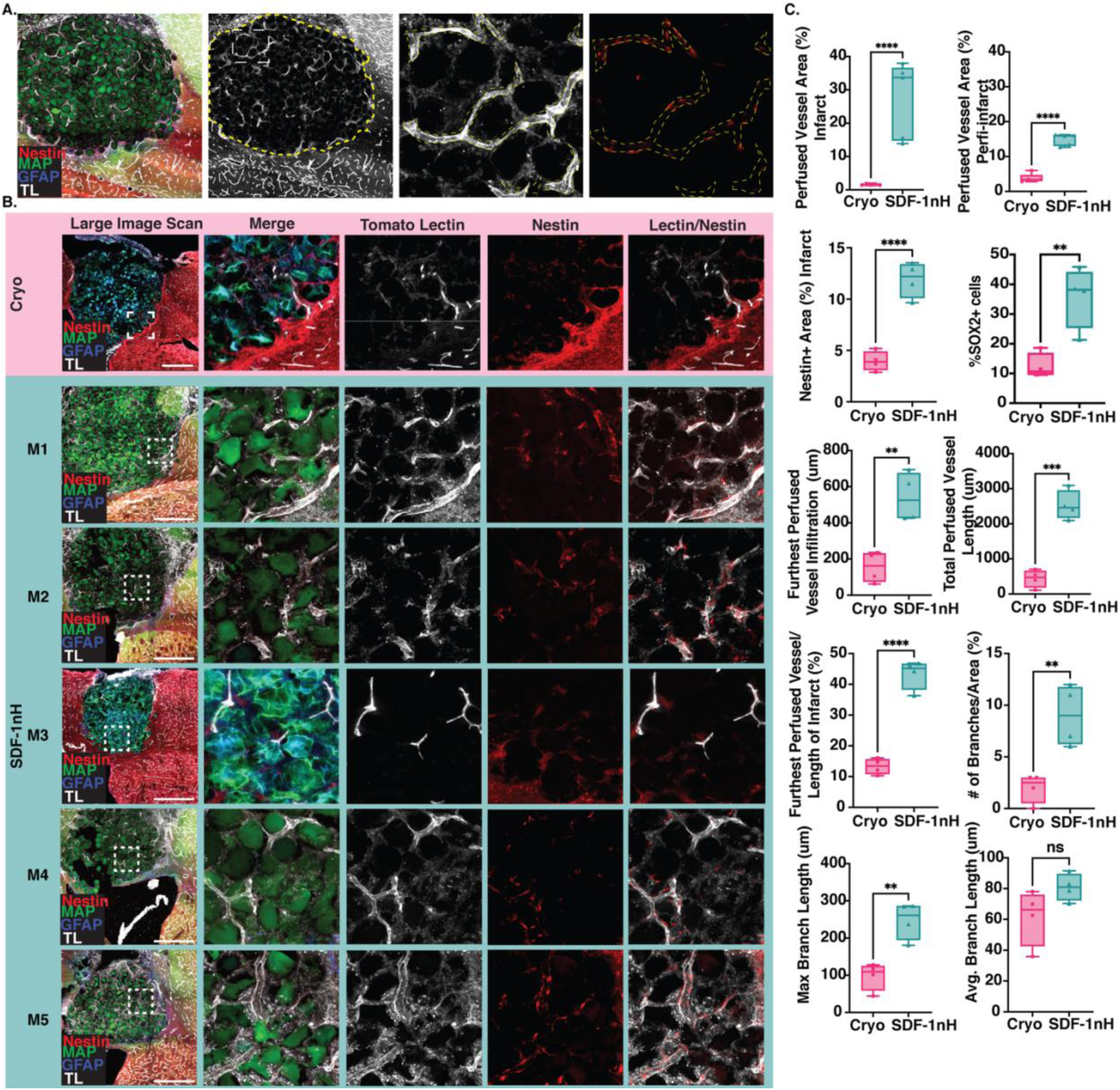
Perfused vasculature characteristics in SDF-1 treated mice 15 days after stroke. A) A 5×5 tiled large image scan of stroke infarct with SDF-1 nH Cryo scaffold (green), Tuj1+ cells (green), TL vessels (white), GFAP+ cells (blue), and Nestin+ cells (red). ROI is dotted in yellow. A magnification of vasculature depicts how it was traced for analysis of co-localization of Nestin+ cells. B) 5×5 20x large image scans followed by a single 20x representative images of Cryo scaffold vs Cryo scaffold SDF-1 nH. Sections of all 5 biological replicates from Cryo scaffold SDF-1 nH are shown. Cryo scaffold (green), vessels (white), Nestin (red). Scale bars represent 500 microns (N=5). C) Quantification of vessel architecture and Nestin with vasculature.

Further analysis on the vasculature showed a significant increase in perfused vessel infiltration averaging 500 µm or 45% of the total infarct distance. This demonstrated that the vasculature was not solely localized to the surrounding peri-infarct or due to boundary effects but observed penetrating all the way to the center of the stroke core (**Figure 6c**). Similarly, the total perfused vessel length, number of branches, and max branch length were all significantly increased in the Cryo scaffold SDF-1 nH group (**Figure 6c**). Vasculature in the peri-infarct was also measured and found to be significantly higher than the Cryo scaffold only group as well (**Figure 6c**), indicating the effects of SDF-1nH extended beyond the stroke cavity. It should be noted that although there is a correlation of SOX2+ cells with increase in vasculature, we cannot state that this is a direct result of NPCs. This is because SDF-1 is also known to recruit endothelial progenitor cells (EPCs),^[15, 23]^ which may be present after stroke given the impaired state of the blood brain barrier. However, both EPCs and NPCs are known to attribute to revascularization and neuroprotective effects after stroke.^[15, 29]^ Analysis of Nestin+ cells within the NestinCre mouse brains show that there is a significant increase in Nestin+ area in Cryo scaffold SDF-1 nH treated mice compared to Cryo scaffold alone. Additionally, we see a significant co-localization of the Nestin+ cells along the vasculature within 300 µm of the stroke border of Cryo scaffold SDF-1 treated mice compared to Cryo scaffold alone (**Figure 6c**).

### 2.7 Incorporation of SDF-1 Does Nothing to Further Attenuate Inflammatory Response after Stroke as Seen with Astrocytes, Macrophage, and Microglia Infiltration

Despite the increase in Tuj1+ cells, vessels, and SOX2+ cells, there was no change in the immune response, measured through glial scar and microglia/macrophage infiltration, when treated with SDF-1 nH (**Figure 7**). After stroke, a glial scar is known to form around the infarct to prevent further damage and limit inflammation to the surrounding area. However, this scar makes it difficult for new tissue to penetrate the infarct and repair the dead tissue. We have previously observed a decrease in this scar, increase in astrocyte infiltration, and attenuated astrocyte and microglial/macrophage reactivity following the injection of MAP scaffolds into the stroke core.^[3, 4]^ As such we stained for astrocytes with GFAP to measure scar thickness from the gel border to the corpus callosum and infiltration. Corpus callosum was not quantified in GFAP scar thickness, as it could be separated from the scar though looking at DAPI. No significant difference was observed between Cryo scaffold and Cryo scaffold SDF-1 nH. Following stains for astrocytes, we stained for microglia/macrophages with Iba1. Again, no significant difference was observed between Cryo scaffold and Cryo scaffold SDF-1 nH treated mice 10 days after injection. It is therefore unlikely that the delivery of SDF-1 is working in any mechanism involving astrocytes or microglia/macrophages. Instead, it is possible the mechanism of SDF-1 promoting vasculature is more centered around progenitor cell activation and recruitment, although further investigation to prove this would be required.

**Figure 7:**
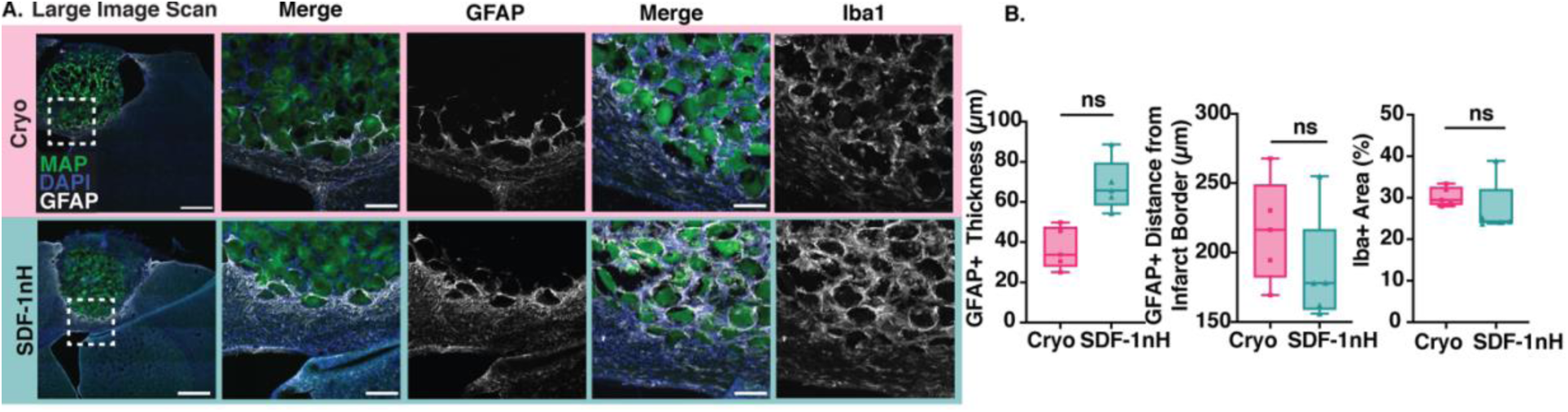
Inflammatory response following the delivery of SDF-1 nanoparticles. A) Representative large scan and magnified images of Cryo scaffold and Cryo scaffold SDF-1 nH treated mice stained for astrocyte (GFAP+, white) or microglial/macrophages (Iba1+, white), DAPI (blue), Nestin (red) and Cryo scaffold (green). Scale bars represent 500 microns in large image scans and 100 microns in other images (N=5). B) Quantification of astrocyte scar thickness, infiltration, and microglia/macrophage percent positive area.

## 3. Conclusion

Here we created 2 novel biomaterials for tissue regeneration after stroke. Cryo scaffolds were shown to have increased porosity compared to MAP scaffolds, but no change in storage or young’s modulus. We demonstrated that the increase in porosity of the Cryo scaffold increased vascularization without the use of added growth factors compared to MAP scaffolds or stroke only. However, despite the increase in Glut1+ vasculature, we observed no difference in immune response compared to MAP scaffolds and a slight decrease in NF200+ cells. To our knowledge, this is the first demonstration of Cryo scaffolds and increased vasculature only 10 days after treatment. However, the resulting vasculature did not penetrate the entire stroke core and remained on the borders of the infarct.

To that end, we sought to creating monodisperse heparin-norbornene nanoparticles that were easy to make, maintained a significant shelf life, and versatile function for binding drugs, small molecules, or growth factors. Both DLS and SEM confirmed heparin nanoparticle synthesis with a narrow PDI. SDF-1 was shown to efficiently bind to the nanoparticles and retain functionality. The SDF-1 nanoparticles were also observed to be retained within the scaffolds, with no release or gradient formation. SDF-1 nanoparticles with Cryo scaffolds led to enhanced Tuj1+ cells and SOX2+ cell recruitment both surrounding and within the stroke infarct. Furthermore, delivery of the covalently bound SDF-1 nanoparticles led to subsequent functional vessel formation within the infarct only 10 days after injection, or 15 days after stroke. Nestin+ cells were seen to surround the vasculature, which could be EPCs. Despite significant changes in cellular response, we did not see any change in astrocyte scar thickness or infiltration or any change in the percent positive area of microglia/macrophages. Further staining could be performed such as Iba1 CD68 costaining to determine if there were differences in reactivity of microglia. ^[51]^ To the best of our knowledge, this is the first-time SDF-1 has been delivered without a gradient and demonstrating enhanced NPC recruitment and retention, Tuj1+ cells, and revascularization throughout the stroke core in only 10 days, all of which are key steps in brain regeneration after stroke.

Although the conventional school of thought has claimed that a gradient is essential for the delivery of SDF-1 to recruit NPCs, our findings demonstrate that this is not the case. Rather, biomaterials with bound growth factors can be just as, or more, effective. We theorize this is due to the endogenous response in the brain where soluble SDF-1 is already released by surrounding cells, such as astrocytes, during the immediate days following stroke. Therefore, the sustained recruitment and/or differentiation of NPCs perhaps requires biomaterials that have maintained growth factor presentation within the scaffold environment. This opens a new avenue for creating biomaterials for the treatment of tissue repair in the CNS depending on the time window of applying the biomaterial. It is likely then that delivery of SDF-1 is affecting vasculature through the recruitment and subsequent cell-cell interaction of progenitor cells. Further staining for CXCR4 presentation may aid in understanding if the SOX2+ cells localized around the vasculature are NPCs or EPCs and give insight on the mechanism of SDF-1 within our scaffold. Successful lineage tracing mouse models such as NestinCreER mice would also be ideal in future experimentation to confirm if NPCs are differentiating into Tuj1+ cells or other cell types, such as astrocytes.

## 4. Experimental section/Methods

### Modification of Hyaluronic Acid with Norbornene

Hyaluronic acid-norbornene (HA-NB) was synthesized as previously described.^[44]^ Briefly, 1.0 g of HA (MW 79,000 Da) (Contipro) was dissolved in 80 mL of 200mM MES buffer (pH 5.5). Once dissolved, 3.1 g of 4-(4,6-Dimethoxy[1.3.5]triazin-2-yl)-4-methylmorpholinium chloride (DMTMM) (MW: 294.74 Da) (TCI America, Portland, OR) (4 molar equivalents) was added to the solution. Once combined the reaction was allowed to stir for 10 min. Then 0.677 mL of 5-Norbornene-2-methanamine (a mixture of isomers) (NMA) (TCI America, Portland, OR) (2 molar equivalents) was added drop-wise into the mixture. The reaction was left to stir at room temperature overnight and then precipitated in 1 L of cold EtOH (200 proof). All precipitates were collected and dissolve in 2 M brine solution and dialyzed against DI water for 30 min, then 1 M brine solution for 30 min. This dialysis process was repeated 3 times and then dialyzed against DI water for 24 hours. The final solution was collected and lyophilized to yield the final product. HA-NB was confirmed by ^1^H-NMR with 33.5% NB functionalization. ^1^H-NMR shifts of pendant norbornenes in D_2_O, δ6.33 and δ6.02 (vinyl protons, endo), and δ6.26 and δ6.23 ppm (vinyl protons, exo), were compared to the HA methyl group δ2.05 ppm to determine functionalization. All equivalents are based on the moles of the HA repeat unit.

### Synthesis of Tetrazine Crosslinker and Alexa Fluors

Tetra-polyethylene glycol-tetrazine (4arm-PEG-Tet) was synthesized as previously described.^[52]^ Briefly, 100 mg of tetra-PEG-SH (MW: 20 000 Da) (NOF America, White Plains, NY) and 15 mg of methyltetrazine-PEG4-maleimide (MW: 514.53 Da) (Kerafast, Boston, MA) were dissolved in 0.5 mL CDCl_3_(maleimide/SH ratio of 1.05), then adding 1 µL of triethylamine (TEA) (0.5 molar equivalent). The mixture was left to stir at room temperature for 4 h. The product was precipitated in 50 mL cold diethyl ether and confirmed by ^1^H-NMR.

Alexa Fluor 488 C2-tetrazine (Alexa488-Tet) was synthesized as previously described.^[44]^ Briefly, 2.8 mg of HS-PEG-SH (MW: 3500 Da) (JenKem Technology USA, Plano, TX) (1 molar equivalent to Alexa Fluor C2 maleimide) and 0.41 mg of methyltetrazine-PEG4-maleimide (MW: 514.53 Da) (Kerafast, Boston, MA) (1 molar equivalent to Alexa Fluor C2 maleimide) were dissolved in 0.17 mL CDCl3. Once dissolved, 0.11 µL of triethylamine (TEA) (0.5 molar equivalent) was added to the mixture. The reaction was left to stir at room temperature overnight. Next, 1 mg of Alexa Fluor 488 C2 maleimide (MW: 1250 Da) (Thermo Fisher Scientific, Waltham, Massachusetts) was dissolved in 0.17 mL CDCl_3_ and then added to the reaction. Once mixed, 0.11 µL of triethylamine (TEA) (0.5 molar equivalent) was then added. The final reaction was left to stir at room temperature overnight. The product was precipitated in 10 mL of cold diethyl ether and dried under vacuum overnight. The resulting product was dissolved in dimethylformamide at 1 mg mL^−1^ and stored at −20 °C until further use.

### Microgel Production and Purification

HA-NB µbeads were produced as previously described.^[44]^ Briefly, a planar flow-focusing microfluidic device was used to create monodisperesed HA-NB microparticles. A 1 mL gel precursor solution was made by dissolving HA-NB in 50mM HEPES pH 7.5, di-thiol MMP sensitive linker peptide (Ac-GCRDGPQGIWGQDRCG-NH2, Genscript) (SH/HA ratio of 14), tris(2-carboxyethyl)phosphine (TCEP) (Sigma-Aldrich, St. Louis, MO) (TCEP/SH ratio of 0.25) and 9.90 mM lithium phenyl(2,4,6-trimethylbenzoyl)phosphinate photo-initiator (LAP) (TCI America, Portland, OR). The final HA-NB in the precursor solution should be at 3.5% (w/v). A 5% (v/v) Span-80 in heavy mineral oil solution was used to pinch the precursor solution at the flow focusing region. The resulting droplets were crosslinked by consistent exposure to UV light (20 mW/cm^2^) off chip using an OmniCure LX500 LED Spot UV curing system controller with a OmnniCure LX500 LED MAX head at 365nm wavelength, at 30% power.

Upon completion of fabrication, the suspension was washed with HEPES buffer under a TC hood until no more oil remained. Following washes, endotoxin tests were performed. Endotoxin concentrations were determined with the Pierce LAL Chromogenic Endotoxin Quantitation Kit (Thermo Fisher Scientific) following the manufacturer’s instructions. Particle endotoxin levels were consistently below 0.2 endotoxin U ml^−1^. Particles were then stored at 4°C until further use.

### Microgel Post-Fabrication Modification

HA-NB microgels or cryogels were modified post-fabrication as previously described under sterile conditions.^[44]^ Excess norbornene groups on the microgels were functionalized with 0.005 mM of Tet-AF 488 in HEPES buffer for at least 1 hour to fluorescently tag the microgels. Prior to their use *in vivo*, microgels were post-modified with RGD (1 mM), IKVAV (0.3 mM) and YIGSR (0.048 mM). The RGD peptide (Ac-RGDSPGERCG-NH2) contained a thiol and thus underwent a thiol-ene click reaction using LAP (9.90 mM) and 20 mW/cm^2^ UV light for 2 min. Upon completion of the functionalization, the suspended µbeads were pelleted by centrifuging at 14,000 rcf for 5 min. IKVAV and YIGSR were synthesized to have a tetrazine group as previously described^[44]^ and therefore reacted via tetrazine-norbornene click reaction. The microgels were washed 3 times with 1xHEPES to get rid of any excess LAP and recovered with the same centrifugation conditions.

### Cryogel fabrication

The following was performed under sterile conditions. Prior to modification, microgels were buffer exchanged by soaking them overnight in 10 mM Tris, 1 mM NaCl, 1 mM CaCl, pH 7.5 buffer. This led to the most consistent formation of crystals within the microgels that left significant defects to create cryogels. After soaking overnight, the microgels were centrifuged to pellet the gel and the buffer discarded. 500 μL of microgels were then placed into a 1.5 μL cryotube and topped off with the previous buffer and mixed thoroughly. The tube was then flash frozen with liquid nitrogen until solid, then lyophilized. The cryogels were then resuspended in HEPES buffer and post-modified as needed. Imaging was performed to ensure croygel formation had occurred. Cryogels were kept at 4°C in HEPES buffer until further use.

### Generation of Scaffolds and Mechanical Testing

Cryo scaffolds annealed as previously described.^[44]^ Briefly, the annealed Tet-MAP gels was measured as the storage modulus (G’) using a plate-to-plate rheometer (Physica MCR, Anton Paar, Ashland, VA). The Tet-MAP scaffolds were formed by combining and mixing HMP with 4arm-PEG-Tet at the desired tetrazine/HA-NB (Tet/HA) ratio of 7. 50 μL of the solution was then placed between 2 sigma-coated (Sigma-Aldrich, St. Louis, MO) slides with 1 mm spacers on either side and fastened using binder clips and incubated for 2 hours at 37 °C unless stated otherwise. This method was used to create triplicate 8 mm by 1 mm height discs of the Tet-MAP scaffolds for rheological testing. A frequency sweep was performed on the hydrogels using a strain of 1% with an angular frequency range of 0.5 to 10 rad/s.

Additional mechanical testing on the scaffolds was performed using a MicroStrain Analyzer. A 2.5N load cell with an 16 mm diameter plate was used at a compression strain rate of 1 mm/min and the hydrogel scaffold was indented 0.2 mm or ∼90% of its total thickness. The Young’s modulus was determined by the slope of the line in the linear region where stress is proportional to strain.

### Modification of Heparin with Norbornene

Heparin sodium salt (14,000 Da, EMD Millipore, Burlington, MA, 375095-500KU) was modified with norbornene similarly to HA. First, 500mg of heparin and 735.5 mg of DMTMM were weighed out into a 20ml glass scintillation vial and dissolved in 10mL of MES buffer. A small stir bar was added, and the reaction was left for to stir for 10 min at room temperature to activate the carboxylic acid. 73.5 μL of NMA was then added to the solution as it stirred vigorously. The reaction was capped and left to stir at room temperature overnight. The next day, NaCl was added to the reaction to obtain a 2M final concentration. The solution was transferred to 6-8K MW dialysis tubing and dialyzed against water for 30 min. Dialysis was changed to 1M NaCl for another 30 min and then to water again for 30 min. This was repeated 3x and finally against water for 24hrs. The solution was then lyophilized, and a H’ NMR was taken to determine percent modification (Figure A11). Proton NMR shifts of pendant norbornenes in D20, δ6.33 and δ6.02 (vinyl protons, endo), and δ6.26 and δ6.23 ppm (vinyl protons, exo), where compared to the heparin methyl group δ2.00 ppm to determine functionalization. All equivalents are based on the moles of the heparin repeat unit. The lyophilized product was stored in a glass scintillation vial at −20°C until further use.

### Nanoparticle Fabrication

Heparin nanoparticles were produced via inverse emulsion. First, a 20% (100mg/500uL) solution of Hep-NB in HEPES (0.3M pH 8.2) was made. Dithiothreitol (DTT, VWR, Cat. No. 97061-338, 0.272 mg) was dissolved in 371 µL of DI water, then 129 µL of 50 mM TCEP. This solution was then added to the Hep-NB to create a final 10% solution of Hep-NB. A 5% surfactant in hexane solution was made with 10mL hexane, 383 µL Span-80, and 123 µL Tween-80. Both the surfactant and Heparin-NB were combined in a scintillation vial and left to stir at 25° C for 15 min. Meanwhile, a 30% solution of ammonium persulfate (APS) (30 mg APS/ 100 µL DI water) was made. After 15 min, the hexane/surfactant solution was sonicated for 15 sec at 10% amplitude. 10 µL of N,N,N’,N’-Tetramethylethylenediamine (TEMED, VWR Cat. No. 97064-684) was added to the heparin-NB solution and then sonicated for another 15 sec at 10% amplitude. Lastly, 20 µL of the 30% solution of Ammonium persulfate (APS, VWR, Cat. No. 97064-592) was added to the mixture and sonicated again for 15 sec at 10% amplitude. The entire reaction was then left to stir overnight at 25°C.

A liquid-liquid extraction was performed for purification. To a separatory funnel, 40 mL of hexane, 10 mL of brine, and the heparin-NB solution was added and extracted. The bottom layer containing the brine and heparin nanoparticles was extracted 5 times with 40 mL of hexane. The nanoparticles were placed into an empty 20 mL glass vial and nitrogen was bubbled through for 5 min. The solution was then transferred to a 100MW dialysis tubing and dialyzed against water for 2 days. After dialysis, the solution was filtered using a 0.2 µm syringe filter and DLS analysis was performed to record the nanoparticle size, averaging about 200 µm (Figure XX). The heparin nanoparticle solution was stored in water at 4°C until further use. Nanoparticles remained stable even after 2 years with no signs of aggregation

### SDF-1 nanoparticle fabrication

SDF-1 nanoparticles were created by covalently binding SDF-1 to heparin-NB nanoparticles. 0.1 µg of heparin/2 µg of SDF-1 in 1xPBS and 10 mM LAP for a total volume of 100 µL was placed into an Eppendorf tube on ice. The solution was exposed to 20mW/cm2 of UV light for 2 min and then filtered via a 100 kDa centricon to remove any excess LAP or unbound SDF-1. First, the centricon was blocked with 1% BSA in 1x PBS with 0.05% Tween-20 for 30 minutes before use. After UV exposure, the SDF-1 nanoparticle solution was spun down in the centricon at max RPMs (14,000 RPM) for 10 min. After each spin, 500 µL of 1xPBS was added and the solution spun down again for a total of 8 times. On the last spin, the centricon was flipped over to retrieve the SDF-1 nanoparticle solution. Each wash was kept for ELISA to determine the concentration of SDF-1 bound to the nanoparticles.

### SDF-1 nanoparticle release assay

MAP scaffolds containing SDF-1 (200ng/ 6 uL) were crosslinked using 4-arm PEG-Tet and incubated at 37°C for 30 min. Once crosslinked, the gels were transferred to a 24 well plate and placed into 600 uL of 1x PBS. At 1 hr, 2 hrs, 3 hrs, 7 hrs, 24hrs, 48hrs, and 10 days, 300 uL of PBS was removed and replaced with fresh PBS. Removed buffer was frozen and kept at −20°C until thawed for SDF-1 ELISA to measure the amount of SDF-1 nanoparticle that was unbound in the scaffold. As expected, no unbound particles were observed even after 10 days.

### SDF-1 ELISAS

SDF-1 ELISA was performed following manufacturer protocol. A standard curve of 2ng/mL to 0 ng/mL was sufficient in calculating unbound SDF-1 from the centricon flowthrough and SDF-1 nanoparticles from the release assay. No solution was diluted for the EILSAS. An average of 30 ng of SDF-1 was calculated to be in all the flowthroughs, leaving 1970 ng of SDF-1 bound to 100 ng of heparin nanoparticles, or 19.7ng of SDF-/heparin-NB nanoparticle. No SDF-1 nanoparticles were detected from the release assay. A positive SDF-1 nanoparticle sample was run to ensure that bound SDF-1 would still illicit accurate ELISA readings (see supplemental data).

### Neural Progenitor Cell (NPCs) Isolation and 2D seeding

*N*PCs were gifted from Dr. Chay Kuo’s lab at Duke University and isolated as previously described.^[44]^ NPCs at P2 were used for *in vitro* experiments and grown in N5 media, with 20 ng/ml EGF/FGF supplementation every other day.

### RT-PCR on NPCs

P2 NPCs in 6-well TC treated plates were exposed to soluble SDF-1 at 0ng (control), 100ng, 200ng, or 400ng for 6hrs, 24hrs, and 48 hrs. Total RNA isolation was performed after exposure following QIAGEN RNeasy Mini Kit (Prod. No. 74106). After isolation, RNA concentration was measured using a NanoQuant plate on the Tecan SPARK® Multimode Microplate Reader following manufacture’s guidelines. Briefly, the plate was cleaned with 70% ethanol and allowed to dry. 2 μL of Rnase free water was used for the blank. The plate was run with the blank and then wiped clean with a lint free wipe before placing samples. 2 μL samples were loaded onto each spot and the plate placed in the carrier. Only RNA samples with an OD 260/OD 280 ratio above 2.0 were used for the following cDNA steps.

iScript cDNA synthesis kit (Bio-Rad, Prod. No 1708891) was used on previously isolated RNA following manufacture’s protocols. Samples were placed in a GeneAmp9700 thermocycler. GAPDH and 18S rRNA were used as internal controls. Taqman primers for GAPDH, 18S rRNA, and CXCR4 (Thermo Scientific, Waltham, MA, Cat. No. 4453320) were used. Detection of GAPDH, 18S rRNA, and CXCR4 were performed using a PCR master mix (TaqMan Universal; Applied Biosystems, Waltham, MA) on a BioRad CFX-PCR system.

### Seeding Neural Progenitor Cells in Microgel Scaffolds

Prior to their use *in vitro*, microgels were post-modified with RGD (1 mM), IKVAV high (0.3 mM) and YIGSR (0.048 mM) as previously described.^[44]^ All microgels were allowed to soak in culture media for a minimum of 2 hours prior to centrifuging them again for “dry” microgels before mixing with NPCs. NPCs were seeded at a density of 10,000 cells/ μL of microgel. After mixing, the mixture of cells and gel were plated and then treated with nothing(control) or nanoparticles only, SDF-1 only (200ngs), or SDF-1 bound nanoparticles (200ngs SDF-1). After 6hrs the samples were stained with live/dead (Calcein AM/EthD-III, Biotum Product No. 30002-T) and imaged on a Nikon Ti Eclipse scanning confocal microscope equipped with a C2 laser with a 20x air objective. The green live fluorescence was used to calculate percentage of cell spreading from max intensity projections using Fiji.

### Photo-thrombotic Stroke

Animal procedures were performed in accordance with the US National Institutes of Health Animal Protection Guidelines and approved by the Chancellor’s Animal Research Committee as well as the Duke Office of Environment Health and Safety. C57BL/6 male mice of 8–12 weeks (Jackson Laboratories, Bar Harbor, ME), were used in the study. A PT stroke was performed as previously described.^[53]^ Briefly, mice were positioned in a stereotaxic instrument and administered Rose Bengal (10 mg/ml, i.p.) and the closed skull at the stereotaxic coordinate 1.8 mm medial/lateral was illuminated with a laser for 10 min at 40mw/cm2 and then again at 2.0 mm medial/lateral for an additional 10 min.

### Hydrogel Injection

Hydrogel injections were performed as previously described.^[53]^ Briefly, 5 days after stroke, mice were injected with 6 μLs of either MAP, Cryo scaffold, or Cryo scaffold + SDF-1 nH. Mice were first anesthetized with 3% isoflurane until unresponsive via toe pinch. They were then positioned in a stereotaxic instrument and their previous incision reopened. The burr hole was located, and the Hamilton syringe containing 10 μL of hydrogel was positioned just above the opening. After setting the z-coordinate to 0, the syringe was lowered 0.780 mm and the hydrogel was injected at a rate of 1 μL/min at a total volume of 6 μL. Then, 5 min after injection completed, the syringe was slowly removed to ensure no hydrogel was displaced. Excess hydrogel from the syringe was cleaned out before prepping for the next mouse. The incision was then closed using vetbond and forceps to ensure the skin did not adhere to the skull. Mice were placed in a clean cage on a heat pad and monitored until they awoke.

### Mouse tissue processing and immunostaining

At 15 days after stroke, mice from all groups were injected with tomato lectin (TL, Vector Laboratories, Newark, CA, Biotinylated, Product No. B-1175-1) which was allowed to circulate for 10 min before being anesthetized and perfused with 1x PBS and 4% PFA. After fixation, their brains were collected, placed into 4% PFA for 2hrs and then transferred to 30% sucrose for 3 days until sectioned, as previously described.^[53]^ Fixed mouse brains were attached to chucks using OCT and 30 μm sections spanning the stroke region were collected on 10 serial positive glass slides that had previously been gelatin coated. Slides were stored at −80°C until used for staining.

Slides were washed with 1x TBS 3×5 min and blocked for 1hr with 1x TBS with 0.03% Triton X and 10% donkey serum blocking buffer. After blocking, slides were stained with primary antibodies as follows: rabbit anti-SOX2 (SOX2, 1:100, Abcam Cambridge, MA, USA, ab97959) for maintained stem cell marker, rabbit anti-beta III Tubulin (Tuj1, 1:100, Abcam Cambridge, MA, USA, ab18207) for neuronal cell marker differentiation, rat anti-glial fibrillary acidic protein (GFAP, 1:300, Invitrogen, 13-0300) for astrocytes, or rabbit anti-microglial (Iba1, 1:250, FUJIFILM Wako, 019-1974, Lot# LEP3218) for labeling microglial and macrophages in blocking buffer at 4°C overnight. Secondary antibodies conjugated to Alexa Fluor 647 (Donkey anti-rabbit, Invitrogen, A31573, Lot# 2083195, or Donkey anti-rat 647 Abcam, Ab150155, Lot# GR3379593-2) were used at 1:500 dilution for 2hr at room temperature. For staining the tomato-lectin, Streptavidin AF647 conjugate (Invitrogen, S32357, Lot# 2352166) was used at 1:500 with the secondary antibodies at rt for 2hr in blocking buffer. DAPI (Sigma-Aldrich, St. Louis, MO, D9542-1MG) at 1:500 was added with the secondary antibodies for staining nuclei. After washing and drying, slides were dehydrated and defat as previously described,^[53]^ before mounting with DPX and a coverslip (Azer Scientific, Morgantown, PA, 22 x 60 mm, No. 1.5, Cat. No. 115226). Stained slides were kept in the dark until image. Stained slides were kept in the dark until imaged. A Nikon Ti Eclipse scanning confocal microscope equipped with a C2 laser with a 20x air objective was used to take 5×5 or 4×5 fluorescent images represented as maximum intensity projections used for analysis.

### Percent Void Volume

Cryogels or Microgels labeled with AF488 were used to make scaffold like those for mechanical testing. After MAP scaffolds or CryoMAP scaffolds were formed, they were then incubated with 70kDa Dextran Tetramethylrhodamine solution for at least 1 hour. After incubation, they were placed on coverslips and imaged at 20x, 2-300 μm z stacks with a step size of 5 μms. ND2 files were saved and later used on IMARIS (Oxford instruments, Morrisville, NC) to create surface renderings and calculate percent void volume by averaging the volume of the cryo/microgel divided by the total volume and the dextran volume divided by the total volume. Technical replicated of each scaffold were used for analysis.

### Imaging Analysis

Fiji/ImageJ was used to analyze mas intensity projection (MIP) tiffs of 20x large image scans. We took 2 brain slices from 1 mouse, 300 to 600 μms apart, to analyze and average to equal 1 n. Brain slices were chosen at the center most part of the stroke to avoid artificially increased boundary effects and allow for characterization cellular infiltration at the center of the stroke core. For microgel characterization and void space analysis, IMARIS (Oxford instruments, Morrisville, NC) software was used to create surface renderings from confocal images of the microgels and the dextran. This data output microgel volume, dextran volume, and total volume, which was used to calculate percent void volume.

### Statistical Analysis

Statistical analysis and plotting were performed using GraphPad Prism 9. Mechanical testing was performed with 3 technical replicate gels. Statistics assumed that gel samples, which were cast independently, were statistically independent from each other. Animal statistical significance was assessed using n of 5 biological replicate data, a 95% confidence interval using a one-way ANOVA with Tukey post-hoc test, unless otherwise noted. All error is reported as the standard deviation of error (SD).

## Supporting Information

Supporting Information is available from the Wiley Online Library or from the author.

## Acknowledgments

We like to acknowledge the National Institutes of Health and the National Institute of Neurological Disorders and Stroke for funding (R01NS079691). This work was performed in part at the Duke University Shared Materials Instrumentation Facility (SMIF), which is supported by the National Science Foundation (award number ECCS-2025064), and with the assistance of Duke’s DLAR breeding core. Author contributions: K.L.W. and T.S. designed the experiments, K.L.W. performed experiments and analyzed the results. L.A.O. performed data analysis and *in vitro* experiments. N.I.J. performed animal experiments. J.D.B. helped in experimental design, data interpretation, and intellectual contributions. N.J.D. for conceptualizing the idea of Cryo scaffolds. K.L.W, L.A.O. and T.S wrote the manuscript, with input from all authors.

