## Supplemental Figures S1 & S2 for "SDF-1 Bound Heparin Nanoparticles Recruit Progenitor Cells for Their Differentiation and Promotion of Angiogenesis After Stroke"

**Supplementary Data**

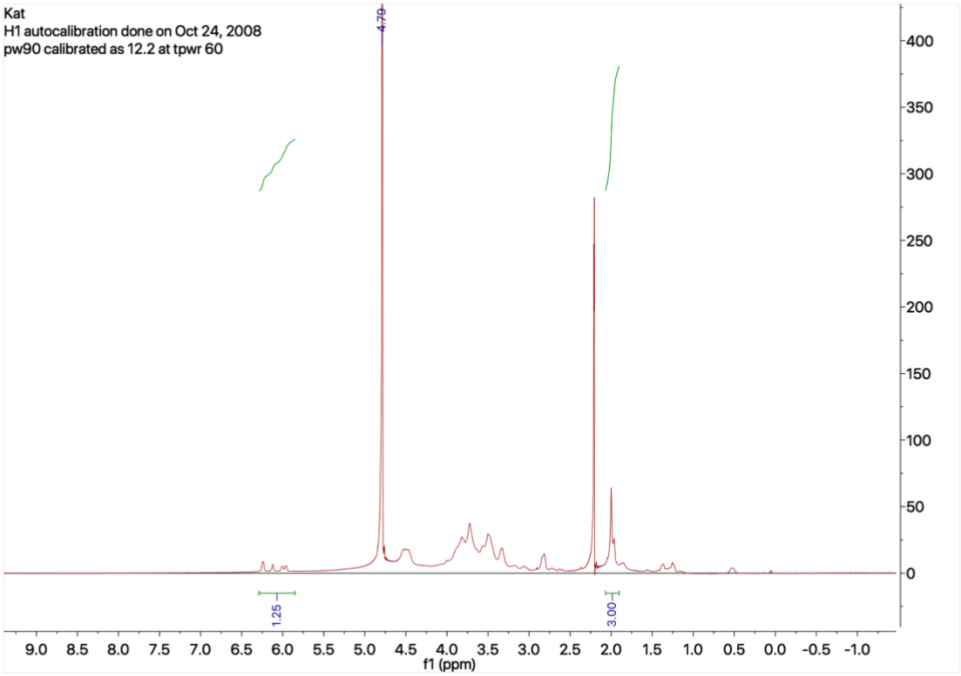


**Figure S1: H'NMR of Heparin-NB**

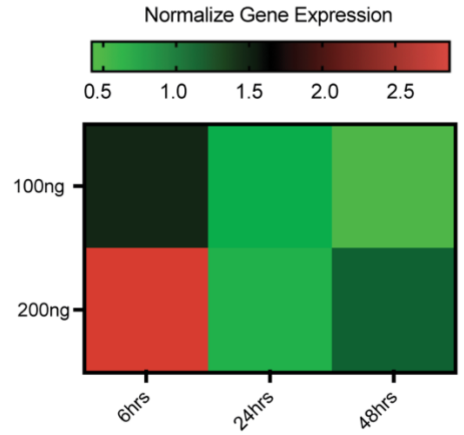


**Figure S2: Heatmap of CXCR4 gene expression over time**
